# P. falciparum PfATP4 Multi-Drug Resistance Resistance to KAE609 (Cipargamin) is Present in Africa

**DOI:** 10.1101/293035

**Authors:** James McCulloch

**Keywords:** Falciparum, Spiroindolones, Cipargamin, KAE609, G223R, Drug Resistance

## Abstract

The PfATP4 (PF3D7 1211900) multi-drug resistance mutation **G223R** is found in Africa by genetically analyzing 2640 worldwide Plasmodium falciparum blood stage isolates (the MalariaGen Pf3k resource). This mutation confers an approximate 8 fold [4] increase in the PfATP4 *IC*_50_ of Spiroindolones (KAE609 & KAE678) [14],[16],[4],[10] and Aminopyrazoles (GNF-Pf4492) [4]. It is postulated that the **G223R** mutation may be a consequence of the drug resistant Southeast Asian Dd2 genotype becoming more dominant in Africa [3]. The presence of this mutation has important policy implications for the eventual general deployment of the Spiroindolone KAE609 (Cipargamin) which is currently undergoing stage 2 clinical trials.

## Introduction

Advances in laboratory automation, process control software and automated biological detectors have allowed malaria researchers to perform high-throughput screening (HTS) of very large chemical libraries [6] [4] to find candidate antimalarial drugs. This HTS screening has been successful in identifying novel anti-malarial drugs including KAE609 [6]. Screens often use whole-cell assays to determine likely candidates. A straightforward method of determining the location and action of the HTS drug candidates is to pressure Plasmodium falciparum (Pf) laboratory strains and map the resultant genetic mutations that confer resistance.

Many resistance conferring mutations of novel HTS candidates have mapped to the transmembrane ATPase sodium pump [15][16] PfATP4 (PF3D7 1211900) indicating that the likely HTS candidate action is the inhibition of this protein. A literature search reveals 54 mutations in PfATP4 conferring resistance. These are tabulated in table 4 for the convenience of the reader. Many of the mutations confer resistance to multiple HTS candidates from different compound classes [16].

The MalariaGen consortium Malaria 3000 (Pf3k) project has processed 2640 genetically sequenced blood stage Pf isolates (128 laboratory, 2512 field) using GATK^1^ best practices into Variant Calling File (VCF) format. This wonderful genetic resource removes the need for the considerable technical and computing infrastructure needed to go from raw short read FASTQ sequence files, via sorted BAM files, to properly generated VCF files^2^. An immediate use of the Pf3k resource is to check for the existence of drug resistance conferring mutations in geographically distinct Pf populations. This was done by Gomes et al [5] for a number of drug target proteins including PfATP4 using *Delly* software (Rausch et al [13]) to analyze Pf3k BAM files and generate variants^3^.

The results tabulated by Gomes et al [5] are different to the results of this research. In particular, Gomes et al [5] do not report the PfATP4 synonymous SNP **G223G** (C669A, 35% of all isolates) characteristic of the Pf strain Dd2 [14] or **G223S** (G667A, 8.9% of all isolates) present in African populations. It should be noted that the G667A and C669A SNPs (GGC*→*AGA) occuring together produce the **G223R** resistance mutation. The probable reason for the difference between Gomes et al [5] and this research appears to be different variant calling methodology. Whereas Gomes et al [5] use *Delly* (Rausch et al [13]) to call variants, this research does not attempt to independently discover variants, but uses the Pf3k VCF files generated by MalariaGen using GATK. The VCF files are then post-processed by `kgl_genome` software developed for this and subsequent Pf genetic research by the author (details in the Methods and Data appendix).

## PfATP4 Mutations in Model Pf Organisms

The MalariaGen Pf3k whole genome sequence (WGS) dataset includes 5 laboratory model Pf reference strains and 121 mixtures and genetic crosses of these model strains (see the MalariaGen ftp website^4^ for more details). Consistent with standard practice, the 3D7^5^ strain is the genetic reference wild type (WT) for this analysis and the MalariaGen BAM/VCF Pf3k generation. The Single Nucleotide Polymorphism (SNP) mutations from these model strains are tabulated below (table 1). The SNP structure of the 121 Pf crosses and mixtures of the 5 model strains is consistent with the pattern of crosses and mixtures of the 5 model strains documented^6^ by MalariaGen. However, the pattern of the SNPs in all isolates (128 laboratory, 2512 field) reflects the geographical bias of the Pf3k field isolates which are only sourced from Southeast Asia and Africa with no South American examples.

**Table 1:** Pf3k Laboratory Strain PfATP4 Mutations

| Pf Strain | Location | E30G | S86N | G223G | Q1081K | G1128R | I1237V |
| --- | --- | --- | --- | --- | --- | --- | --- |
| 3D7 (WT) | Netherlands |  |  |  |  |  |  |
| 7G8 (2) | Brazil | 2 | 2 |  | 2 | 2 | 2 |
| Dd2 (1) | Cambodia |  |  | 1 |  | 1 |  |
| GB4 (1) | Ghana |  |  |  | 1 | 1 |  |
| HB3 (2) | Honduras |  |  |  |  |  |  |
| Mixtures & Crosses (121) |  | 33 | 36 | 30 | 55 | 82 | 34 |
| % of 2640 isolates |  | 1.3% | 1.5% | 35.1% | 36.6% | 56.7% | 3.4% |

## The Geographic Distribution of G223R

The geography of the **G223R** mutation is tabulated below in table 2. The table gives the proportion of the three mutually exclusive mutations of codon 223 in PfATP4 and the synonymous mutation **A229A** (T687A). The Southeast Asian locations that do not have **G223S** or **G223R** (G667A) mutations are not tabulated. The synonymous mutation **G223G** (C669A) is characteristic of the Dd2 model organism [14] and is, as expected, prevalent in Southeast Asia. In particular it is present in 72% of the Ramu, Bangladesh isolates tabulated below. The presence of C669A brings the Pf organism to within 1 SNP of **G223R** with the subsequent addition of G667A. However, only populations with the **G223S** (G667A) mutation have also acquired the **G223R** mutation, strongly suggesting that this mutation is acquired by the subsequent addition of C669A to an *existing* G667A genotype.

**Table 2:** Geographic Locations of G223S and G223R Mutations

| Location | Isolates | G223G<br>C669A | G223R<br>G667A+C669A | G223S<br>G667A | A229A<br>T687A | Linkage DisEq.<br>G667A*T687A |
| --- | --- | --- | --- | --- | --- | --- |
| Nigeria, Ilorin | 5 | 0% (0) | 20% (1) | 0% (0) | 0% (0) |  |
| Guinea, Nzerekore | 100 | 14% (14) | 6% (6) | 16% (16) | 23% (23) | 0.94 |
| Ghana, Kintampo | 68 | 12% (8) | 6% (4) | 24% (16) | 24% (16) | 0.86 |
| Ghana, Navrongo | 549 | 11% (59) | 6% (32) | 22% (119) | 23% (129) | 0.89 |
| Mali, Kolle | 51 | 22% (11) | 4% (2) | 25% (13) | 24% (12) | 0.86 |
| Gambia, Brikama | 65 | 15% (10) | 3% (2) | 12% (8) | 11% (7) | 0.81 |
| Mali, Faladje | 36 | 14% (5) | 3% (1) | 11% (4) | 8% (3) |  |
| CongoDR, Kinshasa | 113 | 18% (20) | 2% (2) | 13% (15) | 12% (14) | 0.89 |
| Senegal, Thies | 133 | 16% (21) | 1% (1) | 12% (16) | 9% (12) | 0.82 |
| Malawi, Chikwawa | 317 | 19% (61) | 1% (2) | 7% (23) | 7% (23) | 0.96 |
| Mali, Bandiagara | 9 | 22% (2) | 0% (0) | 33% (3) | 22% (2) |  |
| Bangladesh, Ramu | 50 | 72% (36) | 0% (0) | 2% (1) | 2% (1) |  |
| Malawi, Zomba | 52 | 10% (5) | 0% (0) | 2% (1) | 2% (1) |  |

The **G223R** mutation may be a consequence of the drug resistant Southeast Asian Dd2 genotype becoming more dominant in Africa [3] with this gene flow adding the Dd2 characteristic mutation **G223G** (C669A) to *existing* **G223S** (G667A) African sub-populations.

We also find very high linkage dis-equilibrium between G667A and T687A (**A229A**). This dis-equilibrium is tabulated as a Pearson correlation (correlations with *p >* 0.01 are not shown). This may be an artifact of an ancestor genotype that has not decayed to linkage equilibrium through recombination because of the proximity of the SNPs. However, there is no similar magnitude of linkage dis-equilibrium with other nearby SNPs such as C669A.

## G223R and Multi-Drug Resistance

Rottmann et al [14] find the **G223R** mutation in one of the Dd2 strains (NITD678-RDd2 clone#1) they pressured with Spiroindolone KAE678 and speculate that the **G223R** mutation may *not* be a result of evolved resistance to KAE678. Manary et al [10] pressured Dd2 strains with KAE678 for 2-4 months and found the **G223R** mutation in the resultant drug resistant strain. Flannery et al [4] show an approximate 8 fold increase in the *IC*_50_ of GNF-Pf4492, KAE609 and KAE678 for G223R mutated Dd2. Spillman and Kirk [16] show that the NITD678-RDd2 clone#1 strain developed by Rottmann et al [14] has an KAE678 *EC*_50_ of 193 *±* 39 nM compared to 21.9 *±* 1.2 nM for the Dd2 wildtype. This is an approximate 9 fold increase and is consistent with results found by Flannery et al [4]. Finally, Jiménez-Díaz et al [7] mention **G223R** as a drug resistant mutation for the Dihydroisoquinolone SJ733 but provide no further detail.

The presence of the **G223R** mutation at moderate but significant levels in Africa has important policy implications for the development and deployment of the new generation of anti-malarial drugs which are active against the ATPase sodium pump PfATP4. One of the most promising of these anti-malarials, KAE609 (Cipargamin) is currently (March 2018) undergoing stage 2 clinical trials. The possibility that the **G223R** mutation may be a result of the drug resistant Southeast Asian Dd2 genotype becoming more dominant in Africa [3] and the very high linkage dis-equilibrium between the SNPs G667A and T687A are interesting topics that merit further research.

## Methods and Data Appendix

This research uses genetically sequenced Plasmodium falciparum isolates from the MalariaGen Pf3K project (release 5, 9 Feb 2016). MalariaGen selected 2640 field (2512) and laboratory (128) isolates from publically available FASTQ short read sequence files. A spreadsheet with accession numbers, location and other useful sample information is available at the ftp website ftp://ngs.sanger.ac.uk/production/pf3k/release_5/. The selected isolates are from Southeast Asia and Africa. MalariaGen processed these isolates into aligned sorted and filtered BAM (compressed SAM) files. The prepared BAM files are then processed by the Broad Institute GATK [12] variant caller to generate VCF files. In order to account for the frequent Multiplicity of Infection (MOI) [11] in Pf blood samples, MalariaGen has assumed a diploid genotype when generating the VCF files.

These VCF files are then post-processed using a C++ software library `kgl_genome` developed for this and subsequent Pf genetic research by the author. This software is available at https://github.com/kellerberrin/KGL_Gene under an MIT license. The associated documentation on GitHub shows how to compile and use this library; the library also has a built-in help function. The `kgl_genome` library uses third party libraries (documented on GitHub) notably the Seqan [2] genome sequence library for advanced sequence comparison and manipulation. Finally, output from `kgl_genome` is examined using tools that will be familar to readers such as Excel and R to produce the final SNP list (no indels were detected in PfATP4). The FASTA and GFF files used by `kgl_genome` were obtained from Plasmodb (http://plasmodb.org) release 35 (1 December 2017).

The identification of SNPs and other variants is a trade-off between sensitivity and specificity; the `kgl_genome` software parses the MalariaGen VCF files and filters variants using the following quality metrics: (1) the Phred-scaled probability that a REF/ALT polymorphism exists, QUAL *≥* 100; (2) the log of the ratio of the probability that a called variant is true over the probability that the called variant is false (log *prob*(*TP*)*/prob*(*FP*)]), VQSLOD^7^ *≥* 2. For each filtered variant, each of the 2640 isolates are individually filtered for depth of coverage, DP *≥* 10 and the Phred-scaled confidence that the isolate genotype assignment is correct, GQ *≥* 20.

After parsing and filtering the VCF file the `kgl_genome` software converts the filtered isolate variant information into an internal variant format and, for each isolate, mutates the reference PfATP4 DNA sequence with the filtered variants obtained for each isolate. This reconstructed mutated DNA sequence is then compared to the reference DNA sequence using the Seqan [2] genome sequence library and a list of variants is generated from the results of this comparison. This indirect method of extracting isolate variant information has the advantage of examining reconstructed mutated DNA using the familiar Needleman Wunsch algorithm and does not assume VCF called variants are independent or mutually exclusive.

This process generates 188 distinct SNP variants from the 2640 isolates. The majority of the distinct SNPs, 110, are found in only 1 or 2 isolates. There are 22 SNPs that are found in *≥* 1% of all isolates and these are tabulated below in table 3.

**Table 3:** All PfATP4 SNPs *≥* 1% frequency in the 2640 Pf3k Isolates.

| Frequency | Chrom. 12 | SNP | Codon | Mutation |
| --- | --- | --- | --- | --- |
| 56.7% (1498) | C529418T | G3382A | GGA→AGA | G1128R |
| 36.6% (966) | G529559T | C3241A | CAA→AAA | Q1081K |
| 35.1% (927) | G532131T | C669A | GGC→GGA | G223G |
| 14.0% (370) | G529707T | C3093A | GGC→GGA | G1031G |
| 9.2% (243) | A532113T | T687A | GCT→GCA | A229A |
| 8.9% (236) | C532133T | G667A | GGC→AGC | G223S |
| 6.8% (180) | A532113C | T687G | GCT→GCG | A229A |
| 5.2% (138) | A529665C | T3135G | AAT→AAG | N1045K |
| 3.5% (93) | T529453C | A3347G | GAT→GGT | D1116G |
| 3.4% (90) | T529091C | A3709G | ATA→GTA | I1237V |
| 2.8% (75) | G531624T | C1176A | ACC→ACA | T392T |
| 2.8% (75) | A529429C | T3371G | CTT→CGT | L1124R |
| 2.2% (59) | C529709T | G3091A | GGC→AGC | G1031S |
| 2.0% (53) | C532133T | G667A | GGC→AGA | } G223R |
| 2.0% (53) | G532131T | C669A | GGC→AGA |  |
| 1.7% (46) | C529454A | G3346T | GAT→TAT | D1116Y |
| 1.5% (40) | C532543T | G257A | AGT→AAT | S86N |
| 1.3% (35) | G530705T | C2095A | CAA→AAA | Q699K |
| 1.3% (34) | T532711C | A89G | GAA→GGA | E30G |
| 1.3% (33) | A531636G | T1164C | GAT→GAC | D388D |
| 1.2% (32) | G531280C | C1520G | ACT→AGT | T507S |
| 1.0% (27) | G529457T | C3343A | CAA→AAA | Q1115K |

## Other Resistance Mutations

A literature search found 54 PfATP4 drug resistance mutations and these are listed in table 4. The 188 distinct SNP variants from the 2640 isolates are examined to see if any occur in the same codon as the tabulated resistance mutations. There are 2 synonymous mutations, P990P (A2970T, CCA*→*CCT, 5 isolates) and Q172Q (A516G, CAA*→*CAG, 1 isolate) that map to resistance codons. However, unlike the synonymous G223G (C669A) SNP, these synonymous SNPs do not increase the probability of the corresponding resistance mutations occurring: P990R (CCA*→*CG*), Q172H (CAA*→*CAC,CAT), Q172K (CAA*→*AAG,AAA).

**Table 4:** Drug Resistance Mutations in PfATP4

| SNP | Compound | Compound Class | Citations |
| --- | --- | --- | --- |
| Q172H | KAE609 | Spiroindolone | [8],[9] |
|  | MMV007275 | Benzamide | [9],[16] |
|  | SJ733 | Dihydroisoquinolones | [7] |
| Q172K | KAE609 | Spiroindolone | [9] |
|  | MMV011567 | Benzamide | [9],[16] |
| V178I | PA21A092 | Pyrazole-amide | [16] |
| A184S | KAE609 | Spiroindolone | [14] |
|  | KAE678 | Spiroindolone | [16],[4],[10] |
| A187V | GNF-Pf4492 | Aminopyrazole | [16],[1] |
| I203M | GNF-Pf4492 | Aminopyrazole | [16],[4] |
|  | KAE609 | Spiroindolone | [14],[4] |
|  | KAE678 | Spiroindolone | [16],[4],[10] |
| V204L | KAE609 | Spiroindolone | [8] |
| A211T | GNF-Pf4492 | Aminopyrazole | [16],[4] |
| G223R | KAE609 | Spiroindolone | [4] |
|  | KAE678 | Spiroindolone | [14],[16],[4],[10] |
|  | GNF-Pf4492 | Aminopyrazole | [4] |
|  | SJ733 | Dihydroisoquinolones | [7] |
| I263V | GNF-Pf4492 | Aminopyrazole | [4] |
|  | KAE609 | Spiroindolone | [14],[4] |
|  | KAE678 | Spiroindolone | [16],[4],[10] |
| S312P | KAE609 | Spiroindolone | [8] |
| L350H | SJ311 | Dihydroisoquinolones | [16] |
|  | SJ733 | Dihydroisoquinolones | [7],[16] |
| L350V | KAE609 | Spiroindolone | [8] |
| A353E | MMV011567 | Benzamide | [16] |
| A353Q | KAE609 | Spiroindolone | [9] |
|  | MMV011567 | Benzamide | [9] |
| N355Y | SJ733 | Dihydroisoquinolones | [7] |
| G358S | SJ733 | Dihydroisoquinolones | [7],[16] |
| I379N | KAE609 | Spiroindolone | [8] |
| I398F | GNF-Pf4492 | Aminopyrazole | [16],[4] |
|  | KAE609 | Spiroindolone | [16],[4],[14],[10] |
|  | KAE678 | Spiroindolone | [10] |
|  | SJ733 | Dihydroisoquinolones | [7] |
| V400A | KAE609 | Spiroindolone | [8] |
| P412L | MMV772 | Undisclosed | [16] |
|  | SJ733 | Dihydroisoquinolones | [7] |
| P412T | SJ733 | Dihydroisoquinolones | [7],[16] |
| V414D | KAE609 | Spiroindolone | [8] |
| V415D | SJ733 | Dihydroisoquinolones | [7],[16] |
| T416N | KAE609 | Spiroindolone | [8] |
| T418N | KAE609 | Spiroindolone | [1],[16],[4],[14],[10] |
|  | SJ733 | Dihydroisoquinolones | [7] |
| A421E | KAE609 | Spiroindolone | [8] |
| P437S | SJ311 | Dihydroisoquinolones | [16],[7] |
| E895K | KAE609 | Spiroindolone | [8] |
| F917L | MMV772 | Undisclosed | [16] |
|  | SJ733 | Dihydroisoquinolones | [7] |
| L928F | SJ733 | Dihydroisoquinolones | [7],[16] |
| L938I | KAE609 | Spiroindolone | [8] |
| P966A | KAE609 | Spiroindolone | [8] |
|  | SJ733 | Dihydroisoquinolones | [7] |
| P966S | SJ247 | Dihydroisoquinolones | [16] |
|  | SJ733 | Dihydroisoquinolones | [7] |
| P966T | SJ279 | Dihydroisoquinolones | [16] |
|  | SJ733 | Dihydroisoquinolones | [7] |
| A967G | KAE609 | Spiroindolone | [8] |
| P990R | GNF-Pf4492 | Aminopyrazole | [1],[16],[4] |
|  | KAE609 | Spiroindolone | [16],[4],[14],[1],[10] |
|  | KAE678 | Spiroindolone | [16],[4],[10] |
|  | SJ733 | Dihydroisoquinolones | [7] |
| A1158V | KAE609 | Spiroindolone | [8] |
| A1207V | KAE609 | Spiroindolone | [8] |
| D1247Y | KAE609 | Spiroindolone | [16],[4],[14],[10] |

## Footnotes

1 The Broad Institute Genome Analysis Toolkit (GATK).

2 For more information see the methods and data appendix

3 *Delly* outputs variants as BCF files but these can be readily converted to VCF format files.

4 ftp://ngs.sanger.ac.uk/production/pf3k/release_5/

5 P. falciparum strain 3D7 was derived from strain NF54 from a patient living in the Netherlands.

6 See the relevant Pf3K documentation at MalariaGen at https://www.malariagen.net/.

7 For a discussion of the VQS, VQSLOD statistics see https://gatkforums.broadinstitute.org/gatk/discussion/39/variant-quality-score-recalibration-vqsr.

